# Neurophysiological mechanisms of immersive virtual reality influence segmental motor reflexes

**DOI:** 10.1101/2025.11.26.690873

**Authors:** Maxim E. Baltin, Anna A. Shulman, Margarita I. Nikulina, Elizaveta A. Vinogradova, Dmitry A. Onishchenko, Diana E. Sabirova, Tatyana V. Baltina

## Abstract

Immersive virtual reality (VR) technologies are being increasingly applied in clinical settings, physiotherapy, and neurorehabilitation due to their potential to modulate neurophysiological functions. However, the mechanistic basis of VR’s effects on human motor systems, especially the impact of emotionally charged VR experiences on segmental spinal reflex excitability, remains unclear. This study aimed to compare the effects of immersive VR stimulation with different emotional tones, the Jendrassik maneuver, and transcranial magnetic stimulation on spinal motor center excitability in healthy adults. H-reflex and M-response amplitudes in the soleus muscle were measured during tibial nerve stimulation under control conditions, during the Jendrassik maneuver, subthreshold magnetic stimulation, and while participants viewed VR videos evoking fear, excitement, or relaxation. The experimental design randomized condition order for each participant. The principal finding was that emotionally salient VR content, particularly scenes inducing fear, produced clear lateralized effects on the spinal motor system. Specifically, inhibitory effects on reflex excitability were observed in the dominant limb, while facilitatory changes occurred in the non-dominant limb. These patterns suggest that immersive VR may differentially engage descending modulatory systems, influencing both sympathoadrenal and corticospinal pathways. In contrast, traditional neuromodulatory interventions did not alter reflex parameters compared to control. The results highlight the unique multimodal influence of immersive VR on sensorimotor regulation and support its incorporation into advanced neurorehabilitation protocols.

## 1. Introduction

The utilization of virtual reality (VR) in healthcare has substantially increased in recent years, finding application in diverse clinical contexts such as physical therapy, rehabilitation, and diagnostic assessment [1]. VR systems are employed for various pathological conditions, including those affecting the central nervous system, where immersive simulators have been shown to improve motor function recovery [2]. Furthermore, VR is increasingly combined with neuromodulation techniques. For instance, combining virtual exercise bikes with transcutaneous spinal cord stimulation in a patient with spinal cord injury resulted in improved motor and sensory functions [3]. In clinical practice, VR is also used for cognitive diagnosis and desensitization in vestibular disorders (for instance, in persistent postural-perceptual dizziness). According to recent research (2020-2025), virtual reality technologies are being actively integrated into the fields of rehabilitation, neuromodulation, and diagnosis, demonstrating significant promise and efficacy in these areas.

Despite its growing clinical adoption, the neurophysiological mechanisms through which immersive visual stimuli affect motor systems remain poorly understood. For example, an analysis of VR’s effect on postural stability demonstrated that somatosensory information and the cerebellum play a more significant role in posture regulation during immersion, suggesting a shift towards a more conscious mechanism for postural correction [4].

Emerging evidence indicates that immersion in stress-inducing VR scenes, such as a virtual fall or being at a height, can reduce the amplitude of the monosynaptic response (H-reflex) in the gastrocnemius muscle, indicating the activation of descending inhibitory influences [5, 6]. However, existing studies primarily address postural control or anxiety, leaving a gap in understanding the effects of a broader range of emotional and sensory stimuli.

It is hypothesized that emotional and sensory stimuli in VR can modulate activity at both the spinal and suprasegmental levels, including the vestibulospinal and reticular pathways [7, 6]. These influences may be implemented through the activation of the sympathetic nervous system, the emotional circuits of the limbic system, and descending corticospinal control, yet a structured model of these interactions has not yet been formed. Furthermore, it remains unclear how stable and lateralized these effects are, and whether the response differs between dominant and non-dominant limbs [8, 9].

Moreover, the degree of involvement of various descending systems – sympathetic, corticospinal, vestibulospinal – and their potential integration during the perception of virtual stimuli remains unclear. In particular, it is unknown how the emotional component of VR (for example, “horror” or “relaxation” video content) modulates spinal motor centers, including presynaptic inhibition of afferents and motoneuronal excitability [7, 9].

Of particular interest is also the comparison of VR interventions with classical methods of modulating spinal reflexes – such as the Jendrassik maneuver, transcranial magnetic stimulation (TMS), motor imagery, and transcutaneous spinal cord stimulation. While the efficacy of these methods in modulating reflex excitability is well-documented, their effects are limited by the short duration and locality of the effect [10, 11]. In contrast, VR’s immersive and multimodal nature offers the potential to engage entire sensorimotor loops and emotional-motivational systems over sustained periods.

Therefore, despite the active clinical use of VR, the nature of the descending control invoked by immersive stimulation remains inadequately defined. A detailed analysis of H- and M-reflexes, as objective electrophysiological markers of segmental activity, can provide crucial insights into these mechanisms.

The objective of this study was to evaluate the state of spinal motor centres under different experimental conditions, including immersion in emotionally distinct VR environments, the Jendrassik maneuver, and transcranial magnetic stimulation.

## 2. Materials and methods

### 2.1 Participants

The study involved 13 healthy volunteers (7 men and 6 women) aged between 20 and 30 years (mean age – 24.1 ± 2.9 years). A thorough clinical examination confirmed the absence of musculoskeletal, neurological, or psychiatric pathologies. All participants were determined to be right-leg dominant. Prior to the experiment, written informed consent was obtained from all individuals. The study included only adult participants; no minors (individuals under 18 years of age) were recruited. The study protocol, including the consent procedure, was reviewed and approved by the Ethics Committee of the Scientific and Technological University ‘Sirius’ and the Local Ethics Committee of Kazan (Volga Region) Federal University. The study was conducted in accordance with the principles of the Helsinki Declaration.

### 2.2 Recording of H- and M-waves

Participants were seated on a treatment couch with their legs hanging freely and without contact with the floor. This specific posture was chosen deliberately because foot support against a surface can provide additional afferent information (support-related afferentation), which is capable of reducing or even suppressing the reflex response by modulating spinal neuronal circuits.

To elicit H- and M-waves, electrical stimulation was applied to the tibial nerve (n. tibialis) in the popliteal fossa. Bipolar recording electrodes were placed over the soleus muscle (m. soleus). The H-reflex recruitment curve was constructed by delivering rectangular pulses of 0.1 ms duration at 10-second intervals. To generate the H-reflex recruitment curve, nerve stimulation was performed using rectangular pulses with a duration of 0.1 ms at 10-second intervals. To obtain the M-response curve, nerve stimulation was carried out using supramaximal impulses. The recording and processing of evoked biopotentials were performed using the basic software package of the Neuro-MVP-4 electromyograph (Neurosoft, Russia). The analyzed parameters included the threshold and area of the H- and M-waves, as well as the H/M amplitude ratio, which served as an index of segmental excitability.

### 2.3 Experimental Protocol

Each participant underwent a series of experimental trials. The sequence of these trials was randomized using a Latin square design to mitigate potential order effects.

Initially, H-reflex and M-wave responses were recorded at rest to establish baseline control values. Subsequent recordings were performed under the following conditions:

- Jendrassik Maneuver. To activate descending pathways and enhance reflex excitability, the participant was instructed to interlock the fingers of both hands in front of the chest and then attempt to pull them apart. The evoked responses were recorded at the moment of hand interlocking [12].
- Transcranial Magnetic Stimulation (TMS). A transcranial magnetic pulse was employed as a conditioning stimulus. The motor evoked potential (MEP) threshold for lower limb muscles was first determined. A 110 mm diameter magnetic coil from the “Neuro-MS/D” stimulator (“Neurosoft,” Ivanovo, Russia) was positioned at the point of intersection of the line from the inion to the glabella and the left and right tragus of the auricle [13]. The MEP threshold was defined as the minimum TMS intensity required to elicit distinct motor responses in 3 out of 5 consecutive trials. The intensity of the subthreshold conditioning TMS was then set individually for each participant at 90% of their resting MEP threshold.
- Virtual Reality (VR) Exposure. Virtual reality was delivered using the HTC Vive system. Three distinct video sequences were selected from the SteamVR library. Recording of H- and M-responses was conducted at the 60-second mark of video playback.

- “Hills” Scenario. A dynamic first-person perspective video of a roller coaster ride was presented through the VR headset, immersing the participant in the simulated environment.
- “Horror” Scenario. A short video featuring frightening visual elements, abrupt sounds, and jump-scare effects was presented to elicit a acute fear response and activate the sympathetic nervous system.
- “Relaxation” Scenario. A video with meditative landscapes, sounds of nature, and soothing music was presented to induce a state of relaxation and reduce emotional arousal. A minimum 10-minute interval was maintained between conditions to allow neurophysiological parameters to return to baseline levels. Consistent ambient lighting, noise levels, and participant posture were ensured throughout the entire procedure.

### 2.4 Statistical Analysis

Data analysis was performed using the Python programming language (version 3.12) with the following libraries: NumPy (2.0.1), SciPy (1.15.1), Matplotlib (3.10.0), and Seaborn (0.13.2). Group comparisons were conducted using either the paired Student’s t-test or the Wilcoxon signed-rank test, depending on data normality and homoscedasticity. No correction for multiple comparisons was applied. Effect sizes were calculated using the absolute value of Cliff’s delta (|δ|). Differences were considered statistically significant at p ≤ 0.05 with an absolute effect size of |δ| ≥ 0.15.

## 3. Results

### 3.1 H-reflex Thresholds

As shown in Figure 2A, significant differences with medium effect sizes (|δ|>0.33) were observed in the right (dominant) leg between experimental conditions. The highest H-reflex threshold was recorded during the scary video condition, and it was significantly higher compared to the Jendrassik maneuver condition (|δ|=0.57, p≤0.05), the relaxing video condition (|δ|=0.49, p≤0.01), and the transcranial magnetic stimulation (TMS) condition (|δ|=0.64, p≤0.01). This indicates an inhibition of spinal reflexes under the influence of fear, likely mediated by the activation of descending inhibitory pathways and an enhancement of presynaptic inhibition.

**Figure 1.**
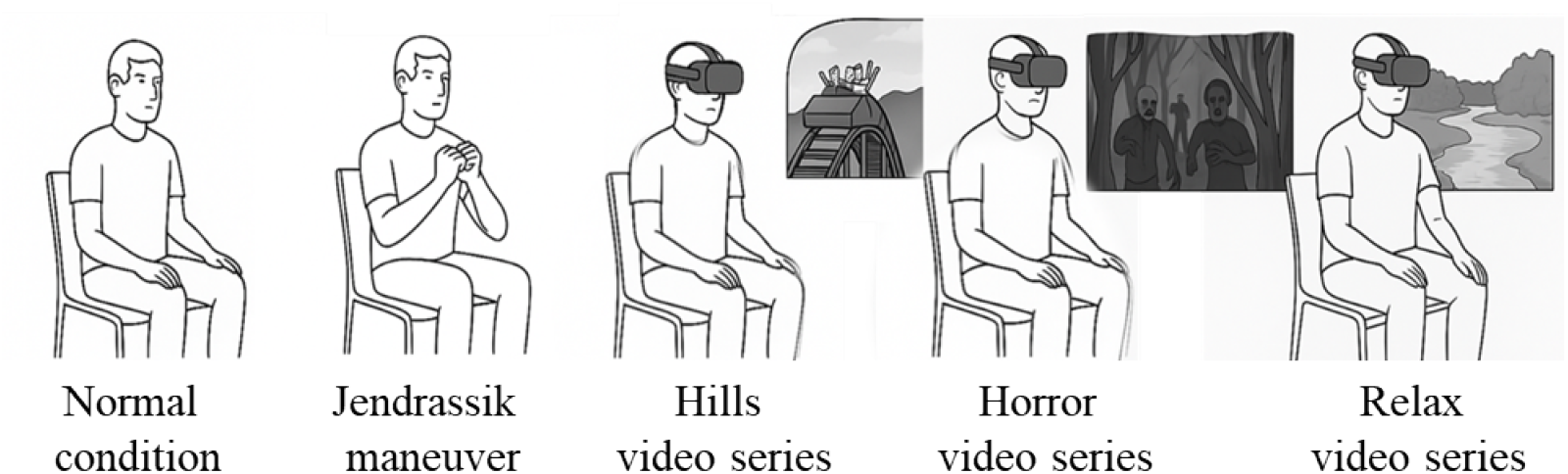
Examples of experimental conditions

**Figure 2.**
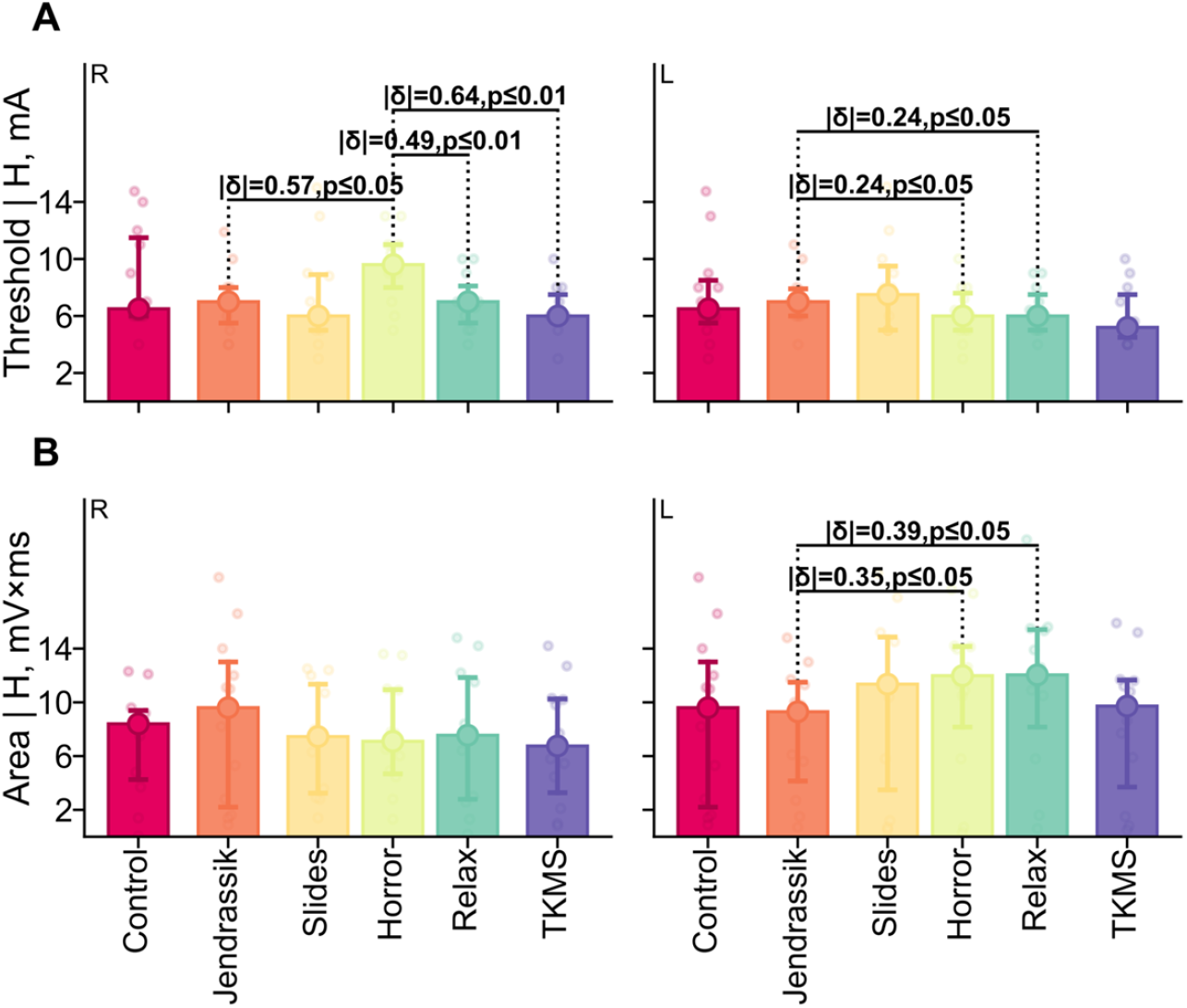
H-response threshold and area under different experimental conditions for the right (R) and left (L) legs. (A) H-response threshold values. (B) H-response area.

In the left (non-dominant) leg, comparisons revealed a significant decrease in the H-reflex threshold during both the scary video (|δ|=0.24, p≤0.05) and the relaxing video (|δ|=0.24, p≤0.05) conditions relative to the Jendrassik maneuver condition, albeit with a small effect size (|δ|>0.15). This may reflect a pronounced facilitation of the reflex arc in response to emotional stimuli, particularly in the non-dominant leg, potentially as part of a lateralized compensatory mechanism.

The obtained results demonstrate that the dominant (right) leg exhibited a significant increase in the monosynaptic reflex (H-reflex) threshold during the presentation of the “horror” video compared to the other experimental conditions. This effect is presumably due to modulation by the central nervous system in response to the perception of a threat, rather than being a consequence of mere sensory stimulation. In contrast to the virtual reality (VR) roller coaster experience, which did not induce significant changes in H-reflexes, only the scary video was accompanied by a significant reflex inhibition in the right limb. This points to a specific role of negatively valenced emotions, which trigger the activation of the sympatho-adrenal axis and a subsequent enhancement of descending presynaptic inhibition of Ia afferents at the spinal level.

Such a response represents a typical defensive modulation aimed at suppressing non-essential motor reflexes in the context of a potential threat [14, 5]. Within the virtual reality (VR) environment, where the stimulus is perceived as more realistic and personally significant, this reaction is particularly pronounced. The activation of the amygdala, hypothalamus, and reticular formation leads to a reduction in the excitability of reflex arcs, but only in the presence of pronounced fear, not mere excitement [15].

Following the “horror” video presentation, an elevation in H-reflex threshold values was registered relative to the control conditions of the Jendrassik maneuver, the relaxing video, and TMS. No significant differences were observed relative to the control for the other conditions. This suggests that the pronounced effect is not so much a facilitation of reflexes induced by TMS, the Jendrassik maneuver, or the relaxing video, but rather a specific inhibitory effect of the video containing frightening, fear-inducing footage on the H-reflex. Thus, the observed significant difference between the “horror” video and the other experimental conditions reflects the induction of emotional inhibition, likely mediated by modulation from the sympathetic and limbic systems [16].

Conversely, in the left (non-dominant) leg, the H-reflex threshold decreased with a small effect size during the scary video condition compared to the other conditions, which may indicate reflex facilitation. This asymmetry may reflect a lateralized response of the central nervous system to emotional stimuli: the right, dominant leg is subject to inhibition, while the left, non-dominant leg experiences heightened reflex activation within a compensatory model [17].

### 3.2 Results for the H-Reflex Area

Figure 2B illustrates the distribution of H-reflex area values under different experimental conditions for the right (R) and left (L) legs. For the right (dominant) limb, no significant differences in the H-reflex area were observed between the experimental conditions (p > 0.05), which may indicate stability in the central regulation of the dominant limb.

In the left (non-dominant) limb, the comparison revealed a significant increase in the H-reflex area during the presentation of the “horror” and “relax” video sequences compared to the Jendrassik maneuver (|δ| = 0.35, p ≤ 0.05 and |δ| = 0.39, p ≤ 0.05, respectively). This may point to a pronounced facilitation of the reflex arc in emotional contexts within the non-dominant limb. The enhancement of the H-reflex area under these conditions reflects a potential activation of descending excitatory influences, including those mediated by autonomic and affective mechanisms.

This suggests that passive emotional states, both anxious (fear) and calm (relaxation), are capable of facilitating spinal reflex activity in the non-dominant limb. Meanwhile, during the Jendrassik maneuver, despite its known effect of reducing presynaptic inhibition, the H-reflex area was smaller in this case, possibly due to differences in the nature of activation. Whereas the Jendrassik maneuver requires voluntary tension of remote muscles, the video stimuli trigger affective and sensory processes (fear, relaxation) that modulate spinal reflexes via descending pathways from the limbic system, amygdala, and brainstem. Such differences between active and passive reflex-modulating influences are supported by studies where emotions (particularly negative ones) enhance the H-reflex by reducing inhibitory interneuronal influences at the spinal level [18, 19].

### 3.3 M-Response Values

Figure 3A shows the distribution of M-wave threshold values under different experimental conditions. The comparison for the dominant (right) leg also revealed no significant effects (p > 0.05). This indicates a relative stability of spinal excitability in the dominant limb amidst changes in emotional and motor background. The absence of a significant variable response might be associated with the function of the right leg as a supporting limb, making it less susceptible to modulation under stress.

**Figure 3.**
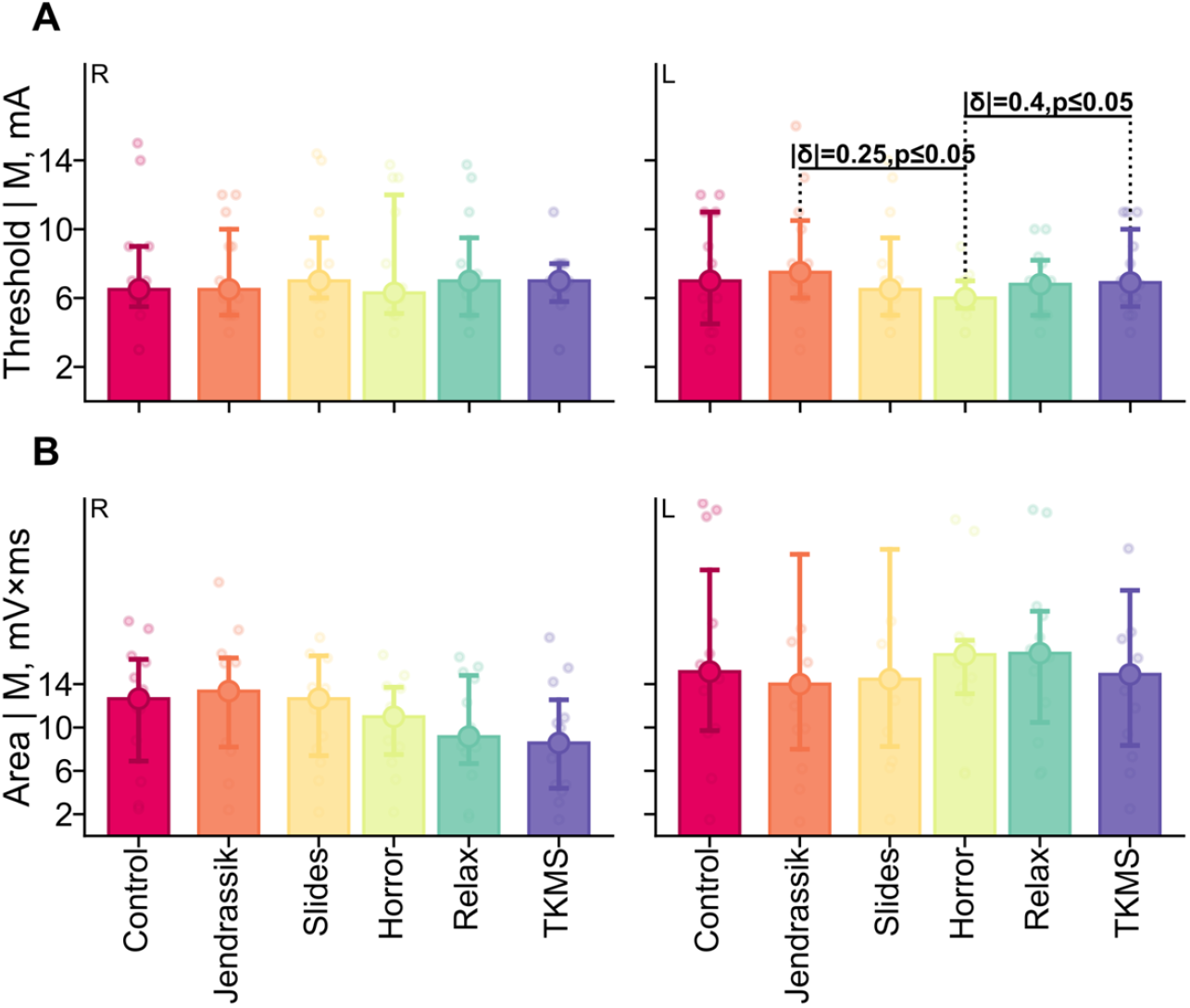
M-response threshold and area under different experimental conditions for the right (R) and left (L) legs. (A) M-response threshold values. (B) M-response area.

**Figure 4.**
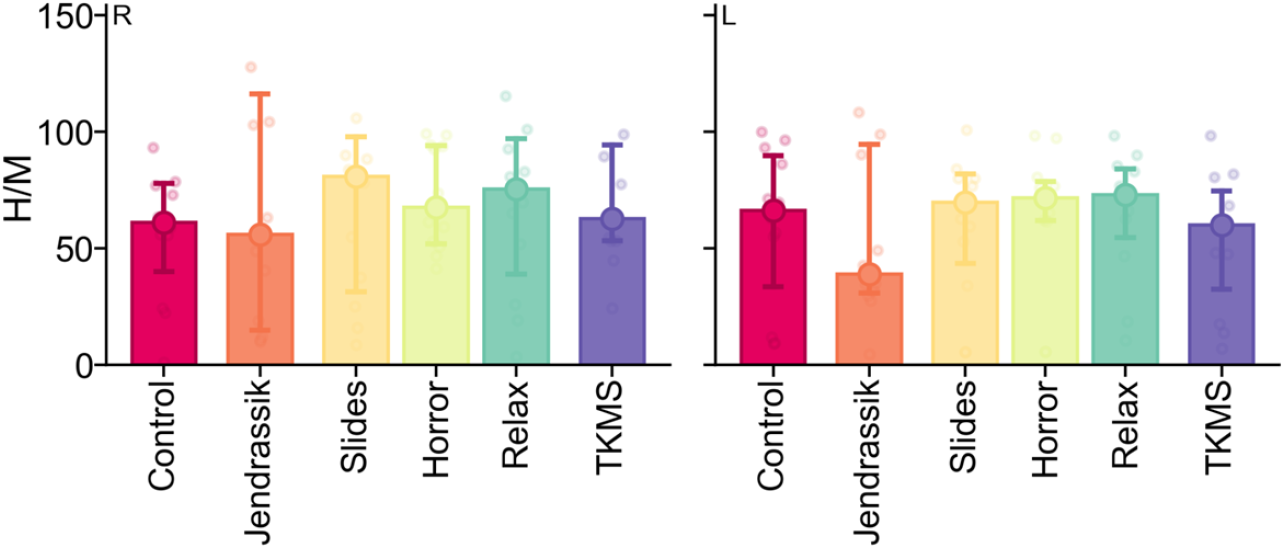
H-response to M-response ratio under different experimental conditions for the right (R) and left (L) legs.

For the data on the left limb, the comparison showed a significant decrease in the M-wave threshold during the presentation of the “horror” video sequence compared to both the Jendrassik maneuver and the TMS group (|δ| = 0.25, p ≤ 0.05 and |δ| = 0.4, p ≤ 0.05, respectively). This indicates that the emotional stress induced by negative VR content can activate motoneuronal circuits and facilitate excitation more effectively than the voluntary Jendrassik test or cortical stimulation (TMS).

Unlike the Jendrassik maneuver, which primarily influences the presynaptic regulation of Ia afferents (and thus has a greater impact on the H-reflex), and unlike TMS, whose effect is limited by stimulation parameters and is primarily associated with central projections, fear induces a global sympatho-adrenal activation. This includes cortico spinal and autonomic influences capable of lowering motoneuron activation thresholds.

### 3.4 The H-Response to M-Response Ratio

The H-response to M-response ratio (H/M ratio) did not demonstrate any significant differences between conditions (p>0.05) in either the right or left leg. The obtained results indicate that the stimuli used did not exert a pronounced influence on the balance between reflex-mediated and direct motor activity.

It is also plausible that changes in the thresholds and amplitudes of the H- and M-responses occurred synchronously (in the same direction), thereby offsetting relative shifts in the H/M ratio. Consequently, this parameter proved to be less sensitive to the intervention compared to the individual parameters.

## 4. Discussion

The present findings demonstrate that emotionally salient VR content simulating a threat (the “horror” video) exerts a pronounced, lateralized influence on segmental reflex activity. In the dominant (right) leg, a reduction in H-reflex excitability was observed, manifested as an increased threshold. These changes are likely mediated by the activation of descending inhibitory pathways, potentially involving sympathetic arousal and enhanced presynaptic inhibition of Ia afferents, a mechanism previously documented in acute psychophysiological stress [22, 16]. Experiments involving virtual simulations of falling report a similar suppression of H-reflex amplitude without changes in background muscle activity [23]. The immersive nature of the VR environment likely amplifies the perceived threat by engaging visual-vestibular and affective channels, potentially recruiting reticulospinal and vestibulospinal pathways in the regulatory response [24].

Conversely, an opposite dynamic was observed in the non-dominant (left) leg, where exposure to the same “horror” video, as well as the relaxation video, led to a decreased H-reflex threshold and an increased H-reflex area. This may indicate compensatory facilitation associated with a lateralized redistribution of neural activity. Emotional stimuli, particularly those with negative valence, are processed predominantly in the right cerebral hemisphere, which can lead to enhanced motor facilitation in the contralateral (left) side of the body [25, 26]. Such lateralized mechanisms have been described in studies of stress responses to acoustic stimuli, accompanied by right-lateralized activation of brainstem and spinal structures [27]. Thus, the observed asymmetry may reflect the involvement of both corticospinal and visceromotor compensatory reactions.

Interestingly, neither the Jendrassik maneuver nor transcranial magnetic stimulation (TMS), despite their known potential to modulate spinal reflexes, produced significant changes compared to the control condition. This may indicate a limited effect of these interventions in the absence of a salient emotional context, or that their effects are transient and subtle compared to the systemic impact of a immersive VR experience. Previous research suggests that the facilitatory effects of motor imagery or TMS are primarily realized when combined with an active task or motor preparation [28, 29].

Furthermore, the H/M ratio failed to show significant differences across conditions. This could be attributed to concurrent, directionally similar changes in both the H-reflex and M-wave components, as well as to the inherent individual variability typical of complex polysynaptic pathways. Nevertheless, the quantitative parameters of the H-reflex and M-wave assessed separately demonstrated high sensitivity to emotional and autonomic modulation, making them preferable metrics for evaluating reflex adaptation.

Collectively, these results confirm that emotionally significant VR exposures can selectively modulate the excitability of segmental motor circuits. The lateralized nature of these effects, alongside the dissociation between passive (emotional) and active (volitional, stimulatory) modulation, underscores the utility of VR as a unique tool for targeting both spinal and supraspinal levels of motor control.

An important limitation of this study is its relatively small sample size, warranting caution in generalizing the findings. Despite the statistically significant differences identified for several parameters, individual variations and demographic factors may have influenced the magnitude of the observed responses. Promising directions for future research include expanding the sample size, stratifying participants based on physiological and psychophysiological profiles, and assessing the sustainability of VR-induced modulation over time and across various clinical populations.

An additional factor that could influence the results is the significant interindividual variability in the effect of the Jendrasik maneuver. In some subjects, this maneuver produced the expected facilitation of the H-reflex due to a decrease in presynaptic inhibition of Ia afferents, while in other participants the facilitatory effect was weak or absent [30]. Such variability may be due to differences in the activation of the fusimotor system, the degree of voluntary muscle tension, and individual characteristics of the descending control of spinal reflexes. Differing or insufficiently pronounced responses to the Jendrasik maneuver could, in turn, influence the evaluation of other experimental conditions, since this test was used as one of the control interventions for comparison with the effects of virtual reality and TMS.

## 5. Conclusions

Exposure to emotionally salient content within an immersive virtual reality environment induces significant changes in neurophysiological parameters, indicative of activated descending regulatory mechanisms. These include sympatho-adrenal and corticospinal modulation, resulting in a lateralized redistribution of spinal excitability.

In contrast, the Jendrassik maneuver, transcranial magnetic stimulation, and neutral or relaxing VR stimuli did not elicit effects of comparable magnitude. This highlights the unique potency of emotional engagement within a virtual environment as a powerful neuromodulatory factor, capable of influencing spinal reflexes and, potentially, motor regulation, with promising implications for clinical and rehabilitation applications.

